# Cellular and biochemical response to chaperone versus substrate reduction therapies in neuropathic Gaucher disease

**DOI:** 10.1101/2021.02.04.429713

**Authors:** Margarita Ivanova, Julia Dao, Neil Kasaci, Benjamin Adewale, Shaista Nazari, Lauren Noll, Jacqueline Fikry, Armaghan Hafez Sanati, Ozlem Goker-Alpan

## Abstract

Gaucher disease (GD) is caused by the deficiency of the lysosomal membrane enzyme glucocerebrosidase (GCase), and the subsequent accumulation of its substrate, glucosylceramide substrate (GC). Mostly missense mutations of the glucocerebrosidase gene (*GBA*) lead to GCase misfolding and inhibiting the lysosome’s proper trafficking. The accumulated GC leads to lysosomal dysfunction and impairs the autophagy pathway.

GD types 2 and 3 (GD2-3), or the neuronopathic forms, affect not only the Central Nervous System (CNS) but also have severe systemic involvement and progressive bone disease. Enzyme replacement therapy (ERT) successfully treats the hematologic manifestations; however, due to the lack of equal distribution of the recombinant enzyme in different organs, it has no impact on the nervous system and has minimal effect on bone involvement. Small molecules have the potential for better tissue distribution. Ambroxol (AMB) is a pharmacologic chaperone that partially recovers the mutated GCase activity and crosses the blood-brain barrier. Eliglustat (EGT) works by inhibiting UDP-glucosylceramide synthase, an enzyme that catalyzes the GC biosynthesis, reducing a GC influx load into the lysosome. Substrate reduction therapy (SRT) using EGT is associated with improvement in GD bone marrow burden score and bone mineral density.

The effects of EGT and ABX on GCase activity and autophagy-lysosomal pathway (ALP) were assessed in primary cell lines derived from patients with GD2-3 and compared to cell lines from healthy controls. While both compounds enhanced GCase activity in control cells, an individualized response was observed in cells from patients with GD2-3 that varied with *GBA* mutations. EGT and AMB enhanced the formation of lysosomal/ late endosomal compartments and autophagy, and this effect was independent of *GBA* mutations. Both AMB and EGT increased mitochondrial mass and density in GD2-3 fibroblasts, suggesting enhancement of the mitochondrial function by activating the mitochondrial membrane potential.

These results suggest that EGT and ABX may have different molecular mechanisms of action, but both enhance GCase activity, improve autophagy-lysosome dynamics and mitochondrial functions.

## Introduction

Gaucher disease (GD) (OMIM 23080, 231000, 231005), the most common lysosomal storage disorder (LSD), is caused by pathologic *GBA* variants (OMIM 606463), resulting in the deficiency of a lysosomal membrane enzyme glucocerebrosidase (GCase) (EC 3.2.1.45). The *GBA* mutations lead to misfolding of GBA in the endoplasmic reticulum, inhibition of proper trafficking and targeting to the lysosomes, and as a result, the deficient enzymatic activity and chronic accumulation of the substrate glucosylceramide (GC) in lysosomes [1]. The major phenotypic presentations of GD are described based on whether the CNS is impacted or not. GD type 1 is the non-neuropathic form, whereas types 2 and 3 (GD2 and GD3) are termed neuropathic GD. GD3 phenotypes are very heterogeneous; however, patients present with horizontal ophthalmoplegia and varying neurological signs, such as progressive myoclonus epilepsy, cerebellar ataxia, cognitive changes, or dementia in some cases [2]. The majority of *GBA* missense variants in patients with GD3 include L444P (L483P) (77%) and D409H (D448H) (7%) [3] [4]. Patients with L444P represent a phenotypically very diverse group with a range of systemic disease severity and neurological involvement [4]. The unique presentation with cardiac involvement, corneal clouding, and hydrocephalus are reported mainly in patients with homozygous for D409H variants [5]. In GD3, the disease onset is before 2 years of age, and in half the cases, often with neurological symptoms. Psychomotor development is affected mostly. Seizures may occur later or manifest as myoclonic epilepsy resistant to antiepileptic drugs. Severe splenomegaly is almost always present and is associated with thrombocytopenia in 60% of cases. Growth retardation (30% of patients) may be the first sign, sometimes associated with cachexia. Lung lesions are sometimes observed, pulmonary infiltration by Gaucher cells, and sometimes due to aspiration. GD type 2 (<5% GD cases) presents in infants aged 3–6 months old with early and severe neurological involvement. The triad consisting of the rigidity of the neck and trunk (opisthotonus), bulbar signs (particularly swallowing abnormalities), and oculomotor paralysis (unilateral or bilateral alternating first then fixed strabismus) is diagnostic of the disease [2]. The mean survival age for GD2 is 11.7 months (range 2–25 months). Before the advent of ERT, children with GD3 succumbed to complications such as portal hypertension and esophageal varices, with significantly reduced lifespans.

ERT is the standard of care in GD for the treatment of systemic symptoms, such as splenomegaly, hepatomegaly, thrombocytopenia, and low platelets [2]. However, ERT is not effective in treating CNS pathology because of the Blood-Brain Barrier (BBB). Other alternative therapy modes using small molecules that may access BBB are SRT and pharmacologic chaperone therapies (PCT).

Glycosphingolipids (GSLs) are involved in a large number of cellular processes, including signal transduction, membrane trafficking, or the formation of cytoskeletal domains. GC is the primary precursor of complex glycosphingolipids, and its synthesis and degradation are the crucial steps for GSLs metabolism. GC is formed by UDP-glucosylceramide synthase (UGCC) in the Golgi apparatus from its precursor ceramide (Fig 1). As an inhibitor of the cytoplasmic enzyme UDP-glucosylceramide synthase (UGCC), EGT is prescribed for type 1 GD patients [6,7]. In clinical trials, EGT has demonstrated efficacy for improvement of systemic disease manifestation, including hepatosplenomegaly, hematologic manifestations, and bone involvement in subjects with GD type 1. [6,8]. EGT therapies are not effective for treating GD’s neuropathic form due to the lack of ability to cross the BBB [3]. A new generation of UGCC inhibitor, venglustat (GZ/SAR402671), could cross the BBB and is in trials for Parkinson, Gaucher, Fabry, and Tay-Sachs diseases [3]. GD and GD-Parkinson mouse studies and cell models currently provide evidence that the related compound (GZ667161) reduces GC’s level in the brain [9].

**Figure 1.**
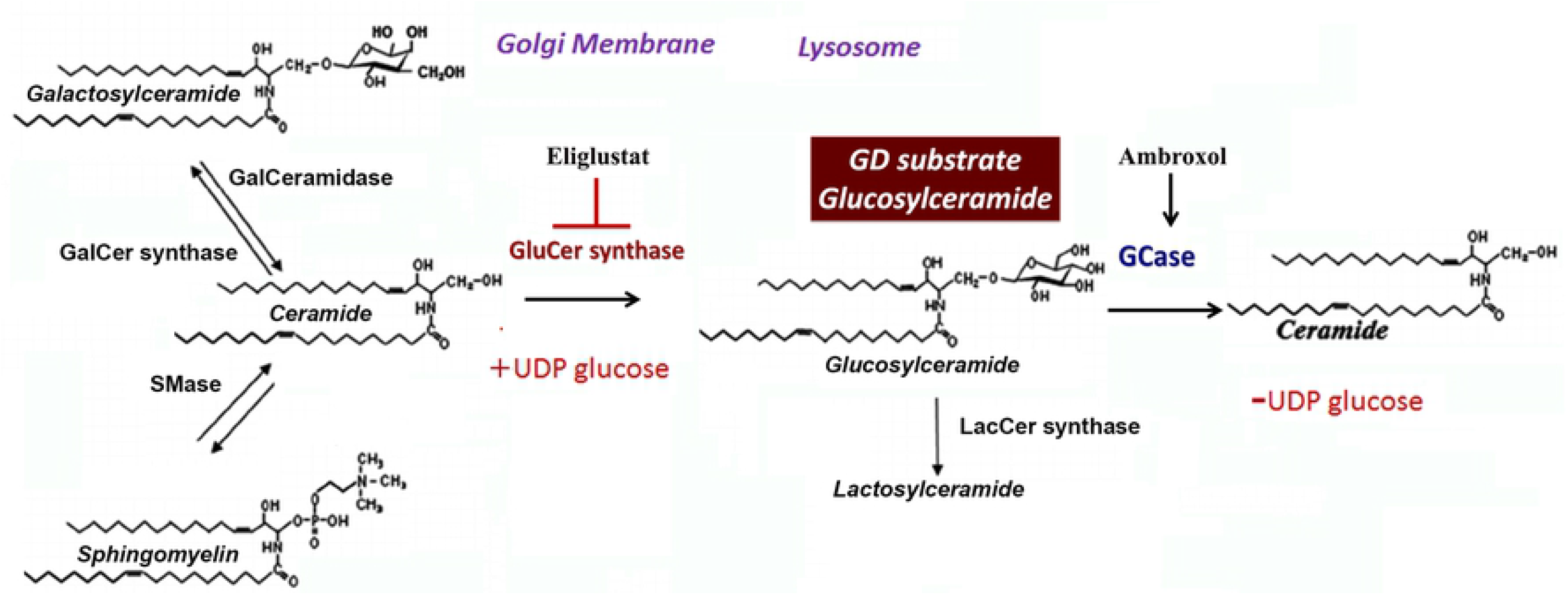
Glucosylceramide (GC) metabolism in Gaucher diseases. Ceramide, glucosylceramide shifts between the Golgi apparatus and lysosomes. UDP-glucosylceramide synthase (GluCer synthase) converts Cer to GC in the ER. GC localized in the intralysosomal membrane is broken down by the GCase enzyme in the presence of SAPC. Eliglustat inhibits GluCer synthase. Ambroxol increases GCase enzyme activity.

Other therapy alternatives for GD using small molecules are enzyme-enhancement (EET) or pharmacologic chaperone (PCT) therapies. PCT is based on small molecules’ ability to fold the misfolded mutant enzyme, deliver an enzyme to the lysosomes, and increase enzymatic activity [10]. The advantage of PCT is that the small molecules can cross the BBB and potentially could treat the neurological symptoms. PCT is a promising approach for GD treatment, especially when the CNS is involved. Recently, an open-label pilot study with ambroxol (AMB) showed promising results in neuronopathic GD patients with N188S, G193W, F213I/RecNciI, and D409H/IVS10^−1^G>A genotypes [11] [12,13]. AMB demonstrated good tolerability while enhancing GCase activity and improving neurological manifestations [14]. AMB interacts with active and non-active sites of enzyme, explaining the mixed type of activation/inhibition and pH-dependent activity [12,15]. Ambroxol demonstrated inhibitory GCase activity at neutral pH and absence of inhibitory effect at the acidic pH of lysosomes. Because different G*BA* mutations have different effects on protein folding, trafficking, and enzyme activity, AMB demonstrates a mutation-dependent chaperoning profile [12,15,16]. We and others recently showed a positive response to AMB in primary cells with N370S/L444P genotype; however, in cells derived from patients with L444P/L444P mutation, there was no uniform GCase activation after AMB treatment [15]. In the present study, we compare the effects of two different therapeutic modalities, SRT and ERT, on primary cells derived from GD2-3 patients. This study aims to investigate the effects of EGT and AMB on GCase enzymatic activity, autophagy-lysosomal pathway, and mitochondrial activity.

## Materials and methods

### Patients

The patients’ demographics and clinical characteristics are presented in **Table 1**. The diagnosis of GD was confirmed by enzymatic activity and molecular analysis. All patients have given written informed consent to collect samples and analyze the data. The clinical protocol was approved by the ethics committees and data protection agencies at all participating sites, protocol # 13-CFCT-07, WIBR protocol #20131424.

**Table 1:** Demographics, clinical and molecular characteristics of subjects with GD.

| # | Gender | Age | Ethnicity | *GBA* genotype | GD Type | GCase residual activity |
| --- | --- | --- | --- | --- | --- | --- |
| 1 | M | 15 | Caucasian | L444P/L444P | GD3 | 4.2% |
| 2 | M | 8 | Caucasian | L444P/L444P | GD3 | 3.8% |
| 3 | F | 1 | Caucasian | L444P/L444P;<br>A456P | GD3 | 0.8% |
| 4 | M | 8 | Asian | L444P/L444P | GD3 | 3.5% |
| 5 | F | 8 | Caucasian | L444P/L444P | GD3 | 4.8% |
| 6 | M | 1 | White | L444P/L444P; RecΔ55, Rec NciI | GD2 | 0.66% |
| 7 | M | 2 | Caucasian | L444P/L444P; R495P/R495P; A456P | GD2 | 2.5% |
| 8 | F | 4 | Hispanic | L444P/L444P D409H/A456P | GD2 | 2.7% |
| **9** | M | <1 | African American | L444P/D409H | GD2-3 | 2% |
| **10\*** | F | 40 | Caucasian | L444P/R493C | GD3 | 4.7% |
| **11\*** | M | 22 | Other | L444P/L444P | GD3 | 10.4% |
| **12\*** | F | 22 | Hispanic | L444P/L444P | GD3 | 9.2% |
| **13\*** | M | 21 | Hispanic | L444P/L444P | GD3 | 9.4% |
| **14\*** | F | 12 | Hispanic | L444P/L444P | GD3 | 8.7% |

• Data from PBMC

### Materials

Eliglustat hemitartrate (EGT) (MCE, MedChem Express, NJ, USA), ambroxol hydrochloride (AMB) (Abcam, Cambridge, UK). Human anti-glucocerebrosidase (GBA) antibodies (GenTex), LAMP1, and LC3A/B antibodies (Cell signaling technology, Danvers, MA, USA). NuPAGE SDS running buffer, bolt 8% Bis-Tris Plus gel, Novex ECL chemiluminescent substrate reagents, sample reducing agents, media 106, low serum growth supplement kit, BCA protein assay kit (Thermo Fisher Scientific, Rockford, IL, USA). Sodium taurocholate hydrate, 4-Methylumbelliferyl β-D-glucopyranoside (Sigma-Aldrich, St. Louis, MO, USA), normocin (InvivoGen, San Diego, CA, USA).

### Isolation and growth of primary skin fibroblasts

Skin biopsies from GD patients were collected to follow the standard procedure. Fibroblast cells were grown in complete M106 media (Life Technologies, Grand Island, NY, USA) as previously described [15]. The primary fibroblasts after passage 5-6 were grown and treated in Dulbecco’s modified Eagle’s media (DMEM) with 10% fetal bovine serum (FBS). Cultures were terminated before passage 10.

### Isolation and purification of peripheral blood monocytes (PBMC)

PBMC were purified from blood samples from GD patients using Lymphoprep™ reagent and SepMate™ tubes (Stemcell Technologies, Vancouver, Canada). Lymphoprep™ was added to the lower compartment of the SepMate tube. Blood was mixed with PBS containing 2% fetal bovine serum (FBS) in a 1:1 ratio and then layered on top of Lymphoprep™ following the manufacturer’s protocol. PBMC cells were cultured in RPMI 1640 media with 5% FBS.

### Differentiation of macrophages from PBMC

Freshly isolated PBMC had been used for macrophage differentiation to follow the procedure described [15]. RPMI 1640 medium with 10% FBS was used to isolate, resuspend, and culture PBMCs. Macrophages differentiation media was RPMI 1640 supplemented with 10% FBS, 1% normocin, 2mM glutamine, 1% Na-pyruvate, 1% non-essential amino acids (NEEA), and 50 ng/ml human recombinant M-CSF (ThermoFisher Scientific, Rockford, IL, USA). After six days of PBMC culture, 100% by volume of fresh complete macrophage differentiation media was added, and two days later, media was replaced with new media. On day ten, macrophages were treated with AMB and EGT indicated in figures intervals and concentrations. GD PBMC normally represents the activated status of macrophages differentiation; some PBMC differentiated into macrophages without external stimulations. In these experiments, PBMC were collected for analysis after treatment, but naturally differentiated macrophages were stained with DALGreen, LysoTracker Red, or MitoTracker Red CMXRos for further evaluation.

### Protein isolation and western blot analysis

Whole-cell extracts (WCE) were prepared in radioimmunoprecipitation (RIPA) buffer. Protein concentrations were determined using the BCA protein assay kit (ThermoFisher Scientific, Rockford, IL, USA). 30 µg of WCE were separated on mini protein TGX stain-free gel and electroblotted using the PVDF transfer membrane (Bio-Rad, Hercules, CA, USA). The ChemiDoc™ MP imaging system (Bio-Rad, Hercules, CA, USA) was used to visualize and quantitate optical density (IOD) for each band. The IODs of bands of interest were normalized to the loading control, beta-actin, and the normalized value of the controls were set to 1 for comparison between separate experiments.

### Measurement of GCase activity

GCase enzymatic activity in cells was carried out using 4-methylumbelliferyl b-D-glucopyranoside. Released 4-methylumbelliferone was measured using a fluorescence plate reader (excitation 360 nm and emission 460 nm) [17,18]. The reaction was started by the addition of 5 or 10 µg of protein into substrates solution in 0.1 M citrated buffer, pH 5.2, supplemented with sodium taurocholate (0.8% w/v). The reaction was terminated by adding 0.4 ml of 0.2 M glycine sodium hydroxide buffer (pH 10.7).

### Measurement of lysosome levels

The LysoTracker Red assay was used to follow the manufacturer’s protocol (LifeTechnology, ThermoFisher, Rockford, IL, USA). LysoTracker Red (50 nM) was added to live cells in the presence of AMB and EGT treatments and stained 30 min. Then, cells were stained with Hoechst, washed 3 times with PBS before analysis. The red fluorescence of LysoTracker was measured in triplicates using a SpectraMax M2 microplate reader with an excitation wavelength: 577 mm; emission wavelength: 590 nm (Molecular Devices, Sunnyvale, CA, USA). Data were normalized to untreated cells. The resulting fluorescence was visualized by fluorescent microscopy (Evos, Hatfield, PA, USA).

### Mitochondrial mass staining

The MitoTracker Red CMXRos mitochondrial kit (ThermoFisher Scientific) was used according to the manufacture protocol to quantify mitochondrial activity upon measure the membrane potential. Nuclear Hoechst dye was used as an index of cell contents. Cells were stained with a fluorescence probe for 30 min and after washed with PBS. MitoTracker Red CMXRos signal was measured in triplicates using SpectraMax M2 microplate reader with excitation/emission 577-590, or the fluorescence signal was visualized by fluorescent microscopy (Evos, Hatfield, PA, USA).

### Mitochondrial membrane potential assay

The mitochondrial membrane potential was determined using the JC-1 Mito-ID membrane Potential Kit (Dojindo Molecular Technologies, Inc). In the energized inner membrane, the mitochondria produced an orange fluorescence signal. If cells exhibit a shift from orange to green fluorescence: mitochondrial function becomes compromised. After five days of treatment with AMB and EGT, fibroblasts were stained with mito-ID membrane potential dye solution in clear-bottom black 96-well tissue culture plates for 30 min. After incubation, cells were washed three times with PBS, and the fluorescence signals were visualized by fluorescent microscopy (Evos Digital microscope, Evos, Hatfield, PA, USA).

### Mitochondrial computational analysis

2D image-based mitochondrial analysis and network characteristics were performed using ImageJ. For network connectivity analysis, the “Skeleton 2D/3D” command was used to calculate the number of branches and branch junctions in the skeletonized network. The analysis tags pixel/voxels in a skeleton image and counts junctions, triple and quadruple points and branches, and program measured junction voxels and endpoints. The voxels are classified as endpoint voxels (if they have less than two neighbors), junction voxels (more than two neighbors). The endpoint voxels are displayed in blue, and junction voxels are displayed in purple. Briefly, fluorescence images of live cells were captured using 40x magnification with large format 2048 × 2048 pixel with the same time exposure and brightness. Selected groups of 2-4 cells were first cropped from the original image to allow analysis on a cell-to-cell basis. The initial contrast of microscope images was enhanced, and residual background pixels were removed follow program algorithm recommendation (ref). The parameters of contrast and background were the same for all images.

### Autophagy staining

DALGreen (Dojindo Molecular Technologies, Inc) was used for the detection of phagosome-lysosome fusion. In several experiments, DALGreen was co-stained with a lysosomal marker, LysoTracker Red. After DALGreen was stained, cells were washed with PBS three times and stained with Hoechst 33342 dye as an index of the nucleus. The resulting fluorescence was visualized by fluorescent microscopy (Evos, Hatfield, PA, USA).

### Glucocerebrosidase immunofluorescence staining

Cells were grown on coverslips and were incubated with 10 μM of AMB and 10 μM of EGT for five days. Cells were then fixed with cold methanol for 5 minutes and washed three times with cold PBS. After blocking with 3% BSA and 0.3% Triton X-100 in PBS for 1 h, primary antibodies GBA (β-glucosidase (A-16), sc-100544, Santa Cruz Biotechnology, Inc, CA, USA) and LAMP1 (D401S, Cell Signalling Technology, MA, USA) were added at a 1:500 dilution for ON +4C^0^. The cells were stained with secondary antibodies labeled with Alexa-Fluor 488 and Alexa-Fluor 555 (ThermoFisher Scientific, Rockford, IL, USA). Cells were incubated with nuclear-DAPI staining. Images were obtained using the Evos^R^ Digital microscope (Evos, Hatfield, PA, USA).

### Cell Viability and cytotoxicity assay

Cell viability was evaluated colorimetrically by the measures the dehydrogenase activity with NADH released in the media using cell counting kit-8 (CCK-8, Dojindo Molecular Technologies, Rockville, MD, USA) according to the manufacturer’s instruction. In brief, cells were seeded on 96-well plates at a density of 50% confluence. Then, the cells were treated with an increasing concentration of AMB and EGT or vehicle control (0.1% DMSO) for 24, 48, 72 h, and 5 days. CCK-8 was added, and absorbance (OD) at 450 nm was detected using the microplate reader (Molecular Devices, Sunnyvale, CA, USA). The IODs were normalized to the untreated control, and the normalized value of the controls was set to 100%.

### ATP assay

The cell titer-Glo luminescent assay was used to measure the ATP level (Promega, Madison, WI, USA). The fibroblasts were plated in 96-well white plates with clear bottom. After AMB and EGT treatments, plates were divided, the half plate was used for CCK-8 assay, and another half of the plate was used to measure ATP. 100 µl of CellTiter-Glo reagent was added directly to the samples, and after 15 min incubation, cells were analyzed by measuring bioluminescence signal in a Genini microplate reader (Molecular Device, San Jose, CA). Samples were run in triplicates.

### LDH release assay

To assess the cytotoxicity potential of the AMB and EGT, the lactate dehydrogenase (LDH) release assay was performed. Control and GD3 fibroblasts were treated with increased concentration of AMB and EGT for 5 days, and supernatants were collected to a new white opaque 96-well plate. After adding the LDH reaction solution (LDH-Glo™ Cytotoxicity Assay, Promega, Madison, WI), the plate was incubated for 30 min. The luminescence signal was read using a Gemini microplate reader (Molecular Device, San Jose, CA).

### Statistical analysis

Statistical analyses were performed using Student’s *t*-test with 2-tailed distribution and 2-sample equal variance or 1-way ANOVA followed by Student-Newman-Keuls using GraphPad Prism (GraphPad, San Diego, CA, USA).

## Results

### EGT, similar to AMB, induces GCase activity

As a GCase chaperone, AMB was demonstrated to increase GCase in cells with N370S/N370S or L444P/N370S *GBA* mutations. However, in cell lines from GD2 or G3 patients with L444P/L444P or L444P in combination with other *GBA* variants, the GCase activity was not uniformly enhanced [12,15,19]. The effect of ETG on GCase activity never was not thoroughly investigated. Thus, EGT and AMB’s effect on GCase enzyme activity was compared in primary fibroblasts derived from GD2 and GD3 patients with different *GBA* mutations (Table 1). Control or GD2-3 fibroblast lines were treated with increasing concentration of AMB and ETG for 5 days, and enzyme activity was measured. EGT increased GCase activity in control fibroblasts in a concentration-dependent manner, similar to AMB (Fig 2A).

**Figure 2.**
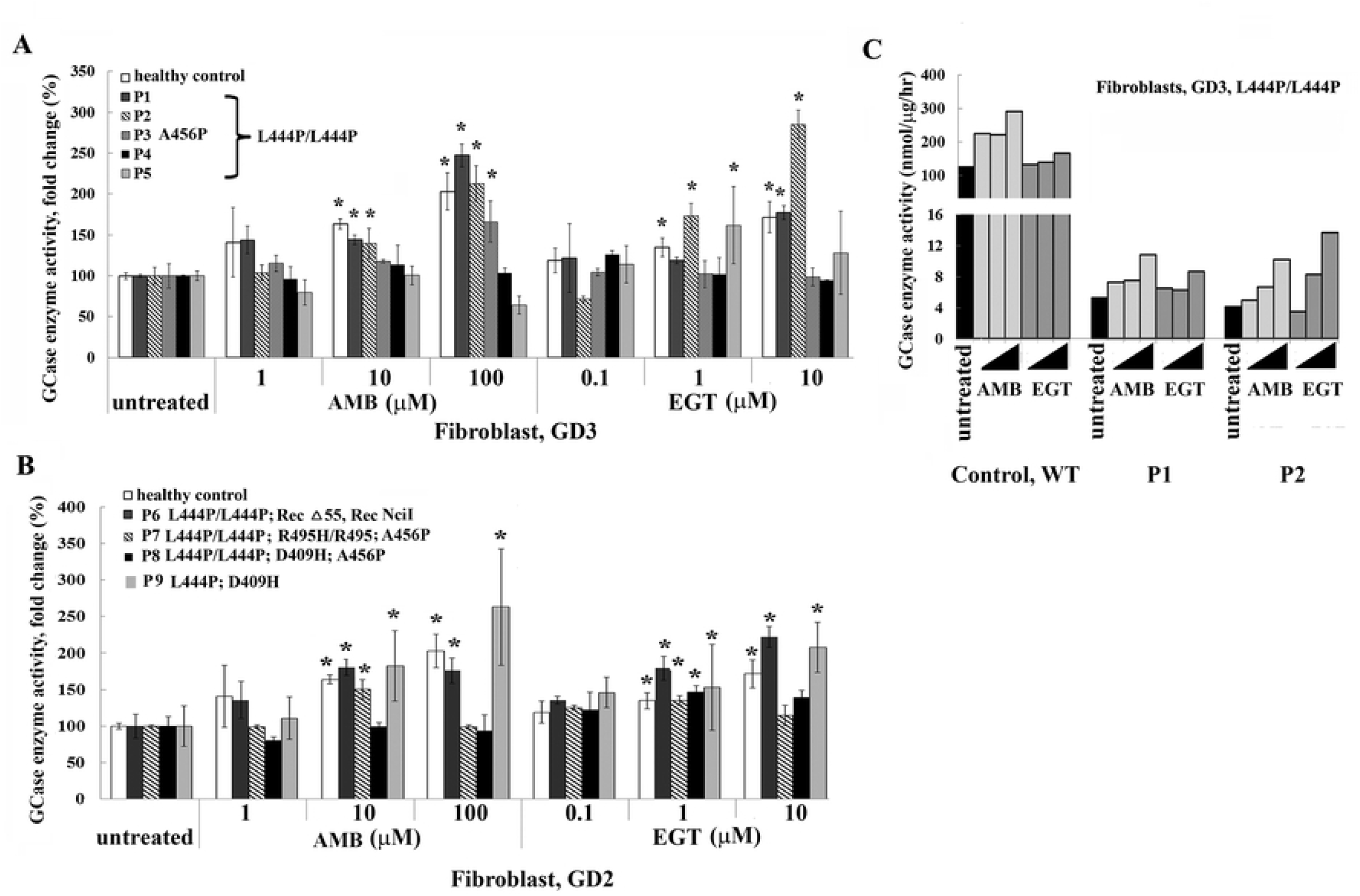
Assessing GCase activity in primary fibroblasts treated with AMB and EGT. **A**. Fibroblasts derived from GD3 patients with the genotypes L444P/L444P and L444P/L444P/A456P were cultured for 5 days in the presence of increasing concentrations of AMB or EGT. Relative GCase enzyme activity was estimated as a fold change towards untreated control. Each bar represents the average +/-STDEV. ∗ p<0.05 compared with an untreated group. **B**. Fibroblasts derived from GD2 with different *GBA* mutations, as indicated, were treated for 5 days in the presence of AMB and EGT. Relative GCase enzyme activity was estimated as a fold change towards untreated control. Each bar represents the average +/-STDEV. ∗ p<0.05 compared with an untreated group. **C**. Comparing GCase enzyme activity estimated as nmol/µg/hr in healthy control fibroblasts and GD3 fibroblasts.

In cells derived from patients with L444P/L444P, there was no uniform GCase activation after AMB and EGT treatments (Fig 2A). AMB and EGT increased GCase activity in P1 and P2 fibroblasts in a concentration-dependent manner (Fig 2A). P3 fibroblasts increased GCase activity in the presence of 100 μM of AMB only, and P5 fibroblasts increased enzyme activity in the presence of 1 μM of EGT. AMB and EGT did not affect GCase activity in P4 fibroblasts (Fig 2A). Overall, 3 out of 5 cell lines showed an elevation of GCase activity in the presence of AMB and EGT (Fig 1A).

GD2 fibroblasts demonstrated personalized responses to AMB and EGT treatments (Fig 2B). P6 and P9 fibroblasts (L444P/L444P; Rec∆55, Rec NciI, and L444P/D409H *GB*A variations) showed increased enzyme activity in the presence of AMB and EGT in a concentration-dependent manner (Fig 2B). P7 fibroblasts (L444P/L444P; R495P/R495P; A456P) showed increased GCase in the presence of 10 μM of AMB and 1 μM of EGT. Only 1 μM of EGT increased GCase in P8 fibroblasts with L444P/L444P; D409H; A456P *GBA* variations (Fig 2B). In absolute terms, the individual levels of GCase in AMB and EGT treated GC cells are still low compare with control (Fig 2C).

PBMC was then collected from five GD3 patients to examine if EGT induced GCase in PBMC and macrophages (S1 Fig). EGT, as an AMB, increased GCase activity in control and GD3 PBMC and macrophages except for P13 GD3 cells (S1A and B Fig).

### Both EGT and AMB improve the lysosomal functions

AMB accelerates the folding and trafficking of GCase to lysosomes and restores the lysosomal functions [15,19,20]. To investigate if EGT mediates the trafficking of GCase to lysosomes, control fibroblasts were treated with both compounds. Staining with anti-GBA (red) and anti-LAMP1 (green) antibodies confirmed that EGT and AMB induce lysosomal localization of GCase (Fig 3A). Then, we compared the effects of EGT and AMB on LAMP1 levels. Western blots showed a significant increase in LAMP1levels in fibroblasts derived from patients with L444P/L444P or L444P/L444P;D409H;A456P *GBA* mutations in response to AMB and EGT treatments (Fig 3B and C).

**Figure 3.**
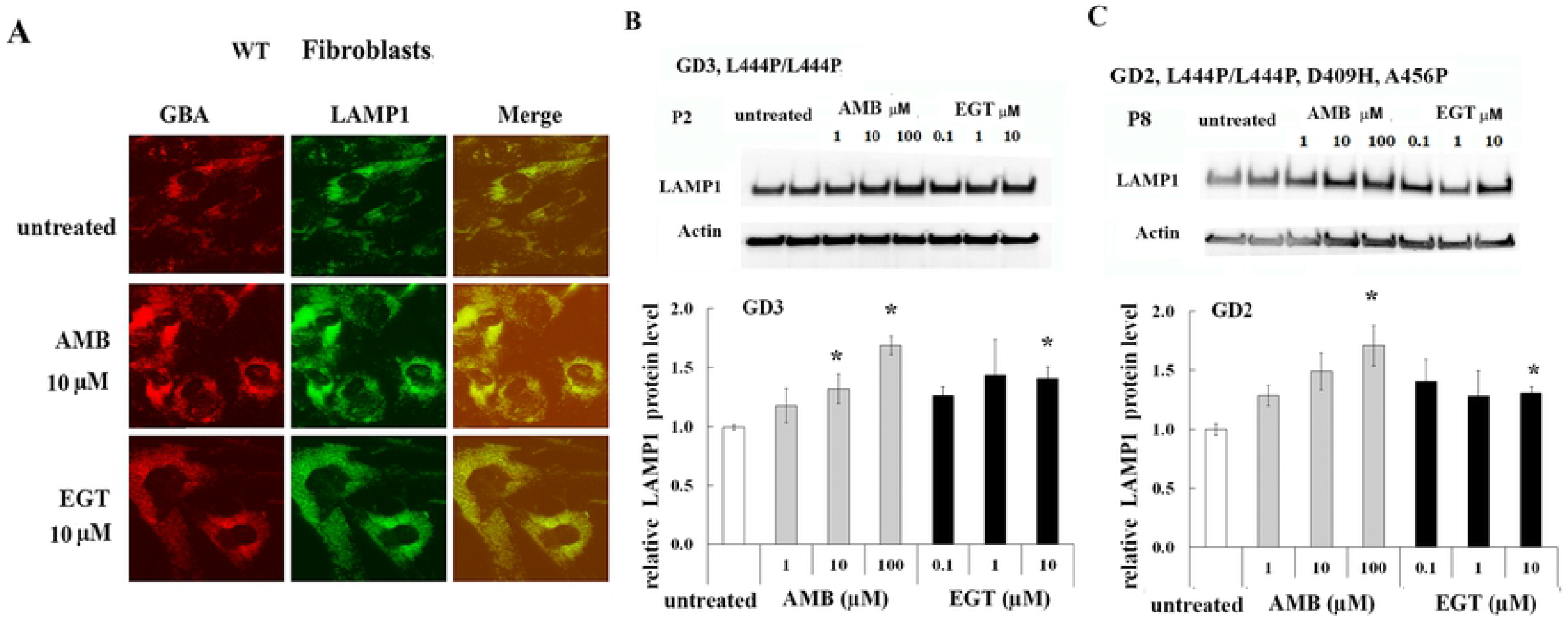
AMB and EGT induced lysosomal trafficking and LAMP-1 level in primary fibroblasts. **A**. Fluorescence microscopic imaging of control fibroblast. As indicated, cells were treated with 10 µM AMB and 10 µM EGT for five days. Each set of three side-by-side images shows anti-GBA (red), anti-LAMP1 (green color) antibodies, and merged images. The yellow color indicates co-localization of GBA and LAMP1 in the lysosome. **B**. Top row represents the western blot of LAMP1 in fibroblasts derived from GD3 patients. Actin is used as the loading control. P1, P2, P4, and P5 (n=4). ∗ p<0.05 compared with an untreated group. **C**. The top: western blot of LAMP1 in GD2 fibroblasts derived from patient 8 with L444P/L444P;D409H;A456P genotype. The bottom: quantification of the relative level of LAMP1/actin from P8. Each bar represents the average +/-STDEV from three independent experiments. ∗ p<0.05 compared with an untreated group.

The LysoTracker Red, an acid-dependent dye, has been used for labeling lysosomal/late endosomal compartments in live cells. Fibroblasts and PBMC were treated with EGT and AMB for five days. Upon treatment with 10, 100 µM of AMB and 1, 10 µM of EGT, the number of acidic vesicles increased in control, GD2, and GD3 fibroblasts (Fig 4). Furthermore, an increase in the LysoTracker Red intensity also was observed in control and GD3 PBMC in the presence of EGT and AMB (S2 Fig). Altogether, the data suggest that activation of lysosomal function is a universal response to EGT and AMB treatments.

**Figure 4.**
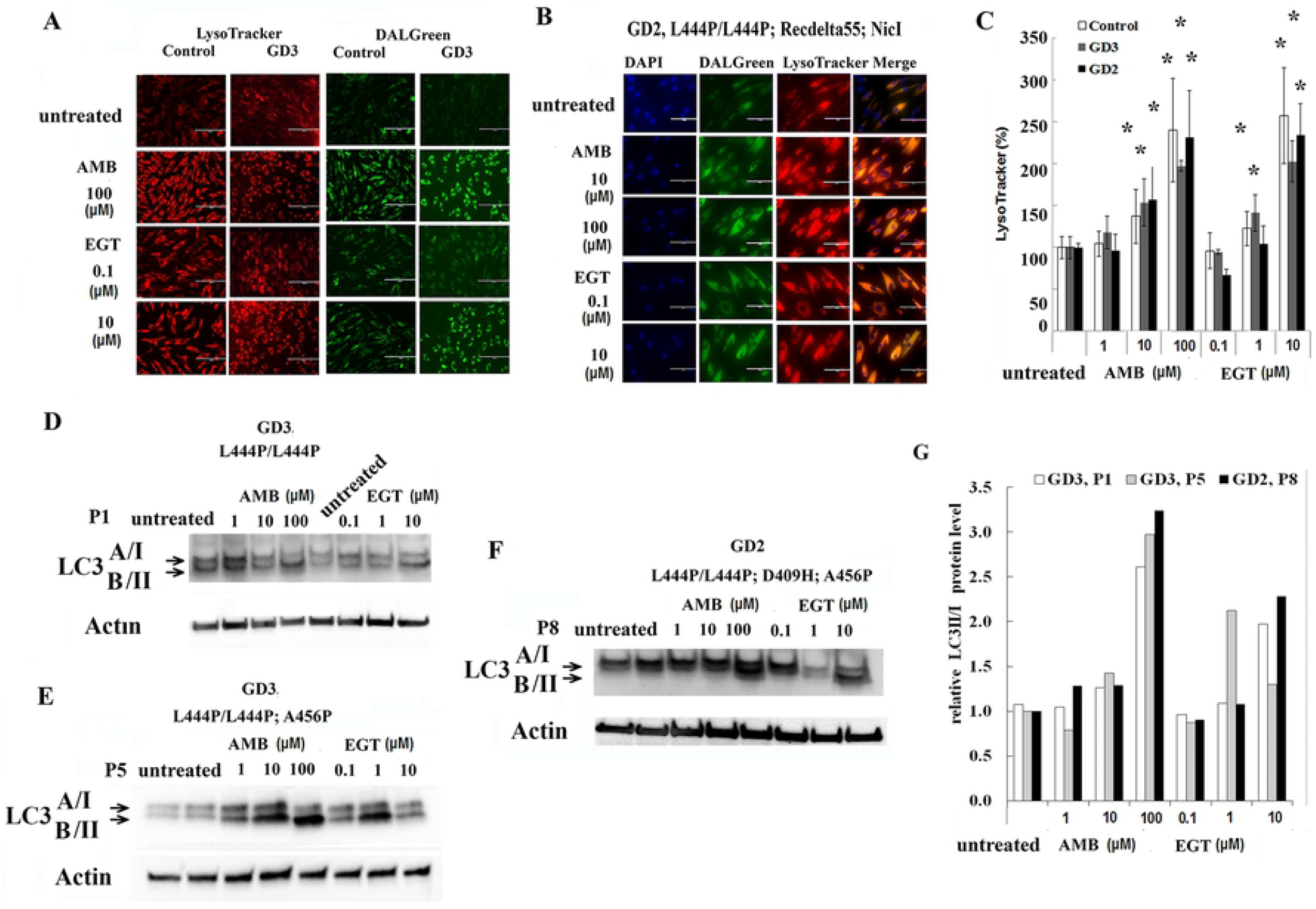
EGT and AMB improve autophagy and lysosomal dynamics. **A**. The lysosome (red) and autophagy (green) fluorescent staining in live control and GD3 (P5) fibroblasts with L444P/L444P *GBA* mutation after AMB and EGT treatments for five days. **B**. Lysosome and autophagy co-localization analysis in GD2 fibroblasts after treatment with AMB and EGT for five days. **C**. Quantification of fluorescence intensity of lysosomes. The signal intensity in untreated cells was set at 100%. The graph indicates the relative intensity value of fluorescence signal related to LysoTracker in control fibroblasts, GD3 fibroblasts with L444P/L444P, and GD2 fibroblasts. Values are expressed as average ± STDEV. **D, E, and F**. Following AMB and EGT treatments, representative western blots showing LC3-I/LC3-II protein expression level in GD3 fibroblast derived from patients: P1 (D), P5 (E), and GD2 patient P8 (F). **G**. Quantification of the relative level of LC3-II to LC3-I.

### EGT and AMB improve autophagy dynamics independent of GCase activation

The accumulation of GC in the lysosomes impairs lysosomal function and inhibits autophagic flux [21,22]. Autophagy and lysosomal staining with DALGreen and LysoTracker Red is used to compare EGT and AMB effects on autophagy-lysosomal function. DALGreen autophagy detection kit is selective for monitoring of late-phase autophagy and autolysosomes [23]. The rate of autolysosomes staining in control, GD3 and GD2 fibroblasts were increased after treatment with 1, 10 µM of EGT and 10, 100 µM of AMB (Fig 4A and B). Merged images confirm autophagosomes’ activation and fusion with lysosomes in GD2 fibroblast (Fig 4B). Like fibroblasts, the autolysosomes staining in PBMC, the staining for auto-lysosomes was significantly increased in control and GD3 cells PBMCPBMC after five days of treatments 10 µM of EGT and 100 µM of AMB (S2 Fig). Merged analysis of autophagosomes with LysoTracker confirmed autophagosome activation and fusion with lysosomes in PBMC (S2 Fig). Autophagy flux marker LC3I/II analysis showed a significantly increased level of LC3-II in GD3 fibroblasts and GD2 fibroblasts with L444P/L444P; D409H; A456P *GBA* mutations after EGT and AMB treatments in a concentration-dependent manner (Fig 4D, E, F, and G). Interestingly, AMB increased autophagy/lysosomal function in P5 fibroblasts without enhancing GCase activity (Fig 4E and G). In summary, AMB and EGT improved autophagy-lysosomal dynamics in primary cells derived from GD2 and GD3 patients.

### EGT and AMB inhibit cell proliferation

To evaluate EGT and AMB cytotoxicity effect, control and GD fibroblasts were treated in the presence of various concentrations of EGT and AMB and then the cell proliferation and viability assay were assessed. The number of viable cells was significantly decreased in control and GD2-3 fibroblasts in a concentration-dependent manner in the presence of AMB and EGT (Fig 5A and B). Analysis of individual cell lines confirmed that the highest concentration of AMB and EGT decreased cell viability in GD fibroblasts derived from patients P5, P6, P7, and P9 (S3 Fig). Moreover, time course treatment verified that the highest concentration of AMB and EGT decreased the number of cells after 24 h treatment (S4 Fig), possibly due to cell proliferation inhibition. Then, we tested the response control, GD2, and GD3 fibroblasts to AMB and EGT exposure by measuring the intracellular ATP level. ATP levels significantly decreased in all cell lines in a dose-dependent manner (Fig 5 and S3 Fig). The ATP/cell viability ratio was analyzed to assess the reason for ATP inhibition: due to reducing the number of cells or due to alteration of mitochondrial function. The ATP/cell viability ratio was increased in control fibroblasts after treatment with 10, 100 μM of AMB, and 1, 10 μM of EGT (Fig 5A). In GD 2-3 cells, 10 μM of AMB and 1 μM of EGT increased ATP/cell viability ratio (Fig 5B). Analysis of individual GD cell lines verified elevation of ATP/cell viability ratio in AMB and EGT treated fibroblasts: P5, P6, and P9 (S3 Fig). AMB, not EGT, displayed the elevation of ATP/cell viability ratio in GD2 fibroblast derived from patient 7 (S3D Fig). The release of the LDH enzyme in media suggests the loss of membrane integrity, the active form of apoptosis or necrosis. The cytotoxic effect of AMB and EGT on control and GD3 fibroblasts, which were treated for five days, were analyzed by LDH assay. The results demonstrated no increase in the level of LDH release in treated cells (S5). In summary, CCK-8, ATP, and LDH results suggest that EGT and AMB inhibit cell proliferation and trigger mitochondrial energy metabolism.

**Figure 5.**
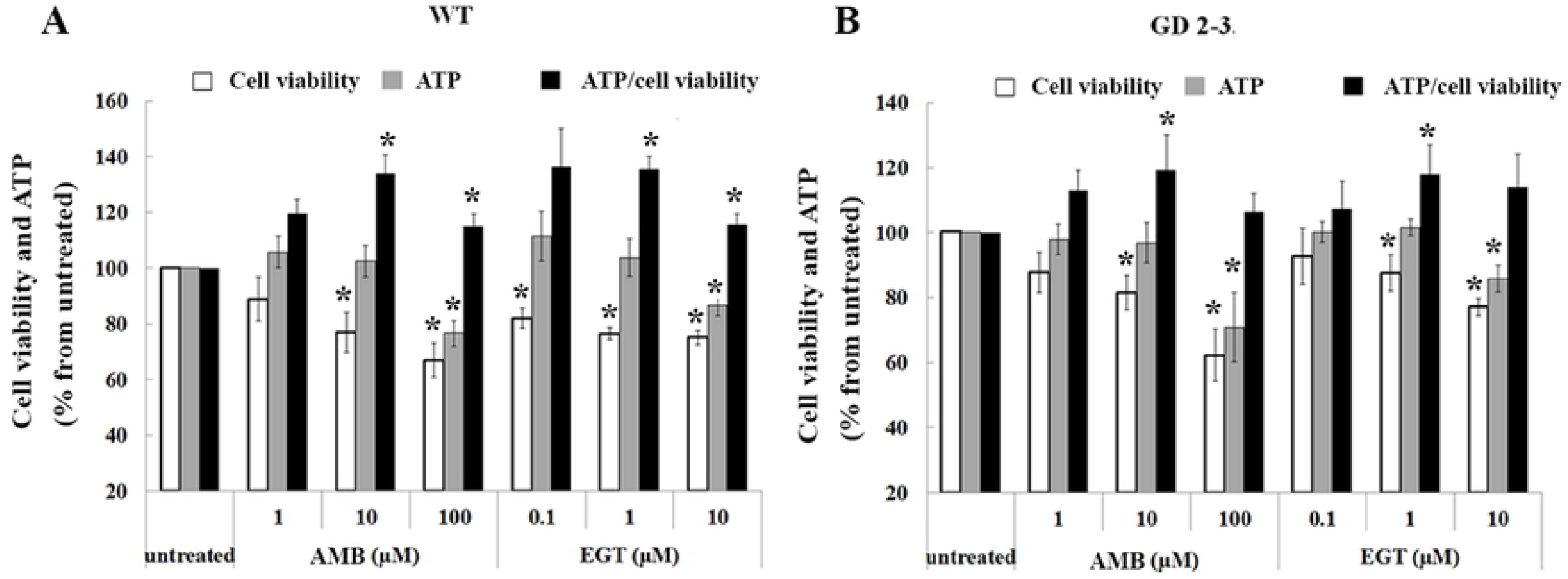
Cell viability and metabolic status in fibroblasts. **A**. Control (WT) fibroblasts were treated with 1, 10 or 100 µM of AMB and 0.1, 1 or 10 µM of EGT for 5 days and were submitted to the CCK-8 cell viability assay and ATP content. The obtain cell counting assay, CCK-8, and ATP results were normalized to the untreated cells. Additionally, the ratio ATP/CCK-8 (cell viability) was estimated. Values are expressed as average ±SEM, n=3. ∗ p<0.05 compared with an untreated group. **B**. Fibroblasts derived from three GD2 patients P6 with L444P/L444P;Rec∆55;RecNCiI, P7 with L444P/L444P;R495P/R495P;A456P *GBA* variations, P9 with L444P/D409H and one GD3 patient with L444P/L444P mutation (P5) were treated with 1, 10, 100 µM of AMB and 0.1, 1,10 µM of EGT for 5 days. The CCK-8 cell viability assay and ATP content were analyzed. The obtain cell counting assay, CCK-8, and ATP results were normalized in relationship to the untreated cells. ATP/CCK-8 (cell viability) ratio was estimated. Values are expressed as average ±SEM, n=4. ∗ p<0.05 compared with an untreated group.

### EGT and AMB enhance mitochondrial metabolism

Because EGT and AMB increased ATP levels in fibroblasts, mitochondrial function was further studied. To visually detect mitochondrial activity in live cells, we used a cell-permeable fluorescent dye, MitoTracker Red. The visual representation of mitochondrial mass and quantitive analysis demonstrated a significant increase in mitochondrial activity in control (Fig 6A) and GD2-3 fibroblasts (Fig 6B) after both treatments in a concentration-dependent manner. Next, we analyzed the mitochondrial density using the skeleton algorithm [24,25]. As shown in Fig 7, the analysis of the 2D images revealed a significant difference in mitochondrial density between untreated and AMB, EGT treated cells. Consistent with increased mitochondrial density, the number of junction voxels and endpoint voxels were increased in cells in the presence of AMB and EGT (Fig 7B).

**Figure 6.**
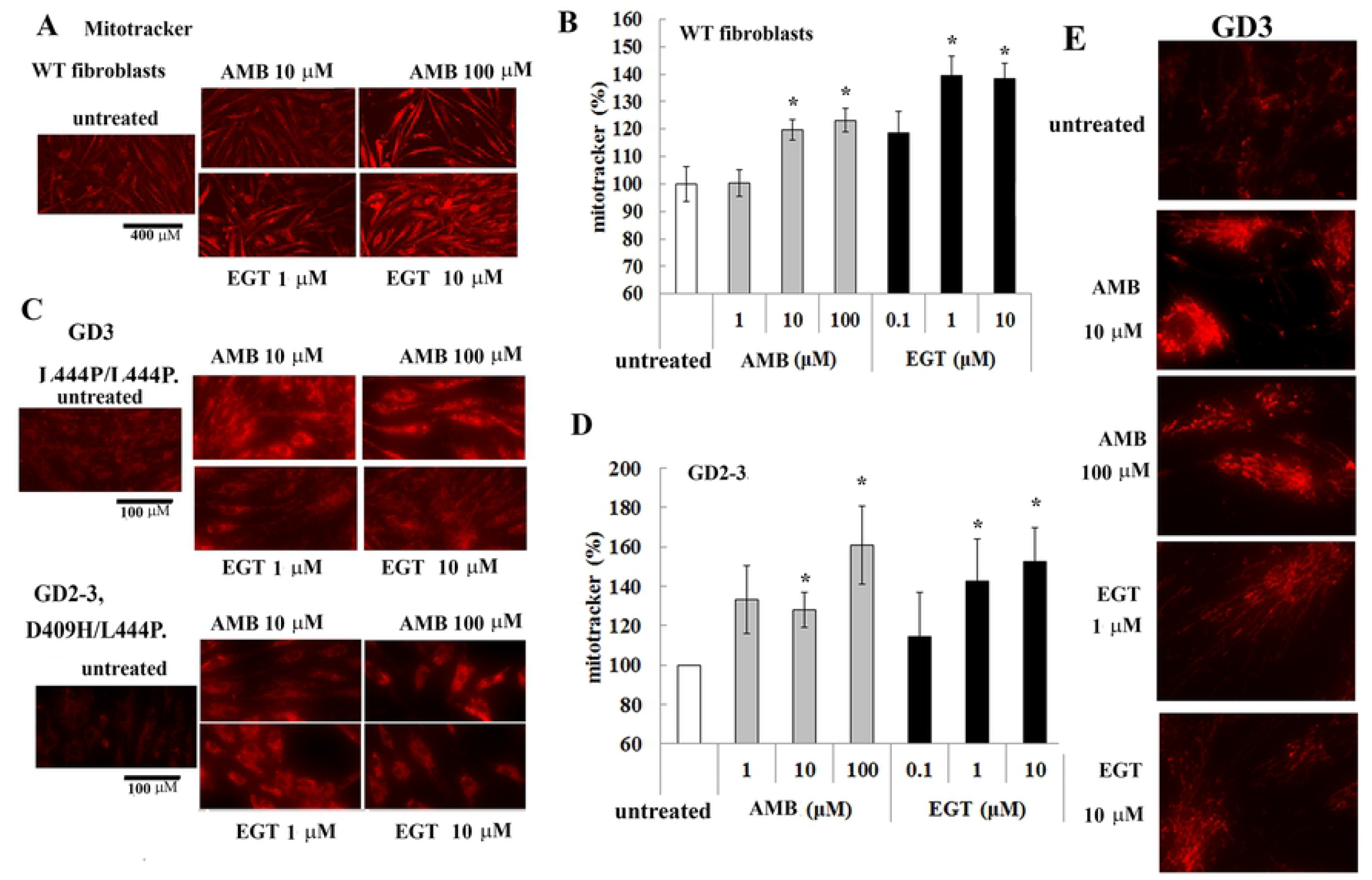
MitoTracker Deep Red staining intensity in AMB and EGT treated fibroblasts. **(A)** The mitochondrial visualization in live control (WT) fibroblasts with AMB and EGT treatment for five days. Scale bar represents 400 µm. **(B)** Quantification of fluorescent intensities of mitochondria. The signal intensity in untreated cells was set at 100%. The graph indicates the relative intensity value of the fluorescence signal related to MitoTracker Red in control fibroblasts. Values are expressed as average ±STDEV. **(C)** The mitochondrial visualization in GD fibroblasts with AMB and EGT treatment for five days. Scale bar represents 100 µm. **(D)** Quantification of fluorescent intensities of mitochondria in GD cells. The signal intensity in untreated cells was set at 100%. Values are expressed as average ±STDEV. ∗ p<0.05 compared with an untreated group. **(E)** The mitochondrial visualization in live GD3 fibroblasts. Representative images were assessed in regards to the degree of mitochondrial branching.

**Figure 7.**
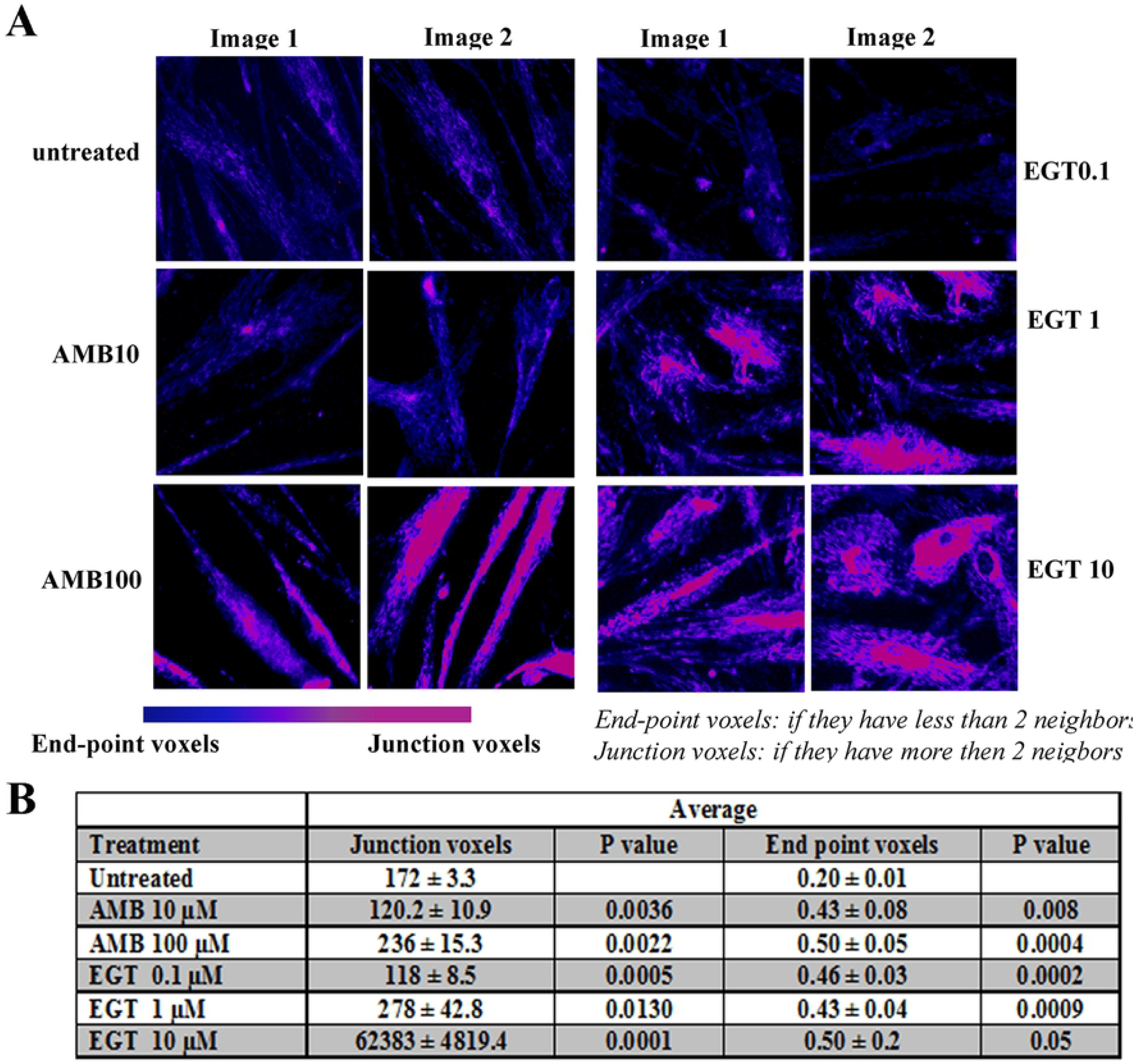
The skeleton algorithm identifies mitochondrial density in sample images from cells. **(A)** Post-processed images of living mitochondria stained with MitoTracker. Cells were treated with AMB and EGT for five days. Two-tone color (blue turn to pink) represents the intensity of density. **(B)** Results of the corresponding mitochondrial assessment using skeleton 2D/3D analysis.

The mitochondrial membrane potential (ΔѰm) is generated by proton pumps (Complexes I, III, and IV), and is an essential component of the ATP process during oxidative phosphorylation. Normally, the levels of ATP and ΔΨm in the cell are kept stable. However, a decrease or rise of ΔΨm may induce. Therefore, we measured ΔѰm using JC-1 dye in GD3 fibroblast and PBMC after treatment with 1, 10, 100 μM of AMB and 0.1, 1 and 10 μM of EGT for 5 days (Fig 8A and B). Healthy mitochondria that have high ΔѰm uptake dye and emit red fluorescence at 590 nm, damaged mitochondria with low ΔѰm emit green fluorescence. JC-1 assay showed that AMB and EGT treatments increased red fluorescence intensity compared with untreated groups indicating activation of hyperpolarization of mitochondria. Surprisingly, 100 µM of AMB treatment increased green and red fluorescence intensity signal, especially in PBMC, indicating the presence of both cytoplasmic JC-1 monomer and mitochondrial J-aggregates in cells (Fig 8).

**Figure 8.**
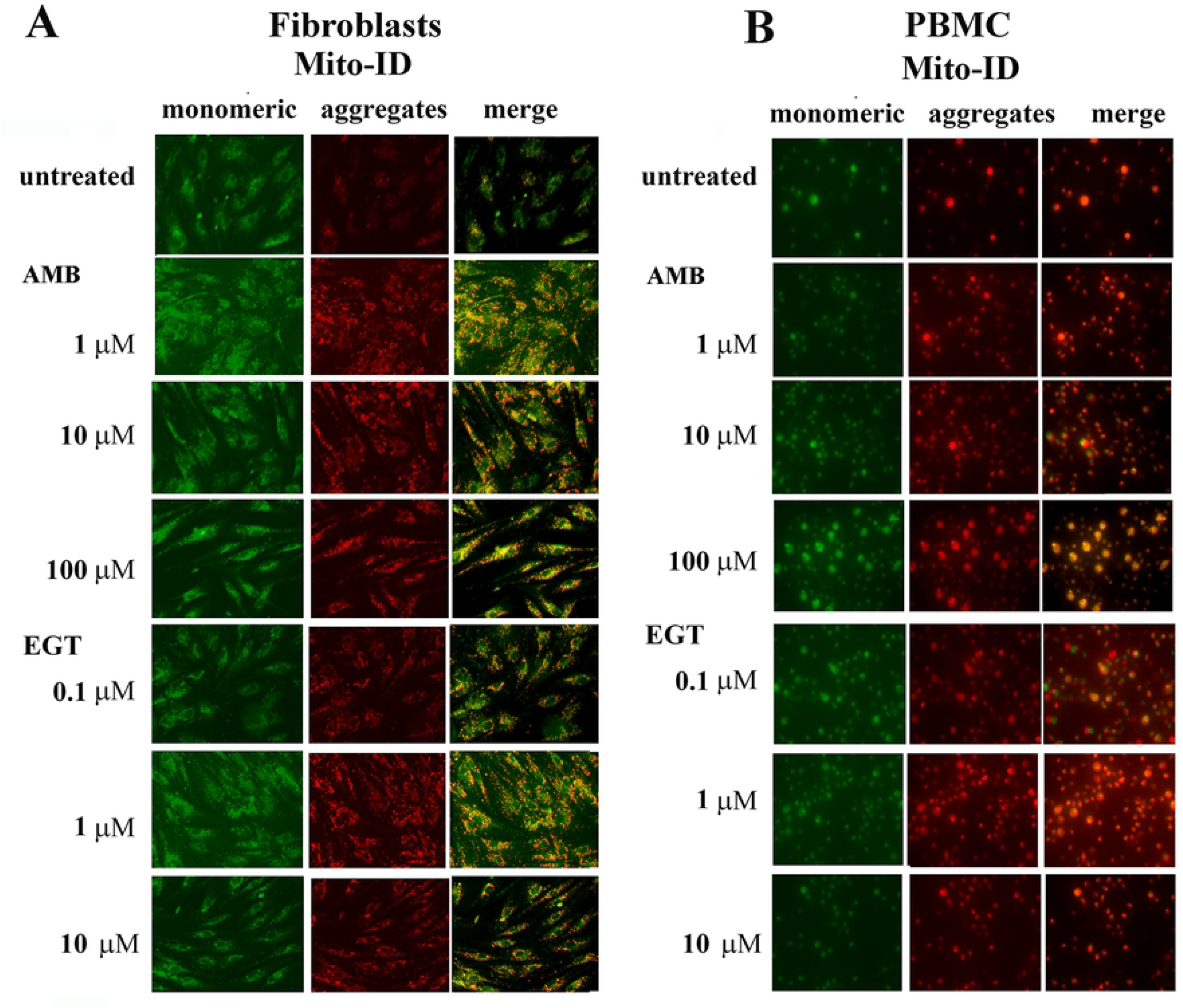
The effects of EGT and AMB on mitochondrial membrane potential GD fibroblasts were treated with increasing concentrations of AMB or EGTfor five days, then were stained with Mito-ID Membrane Potential reagent and visualized by fluorescence microscopy. Green represents mitochondria with low membrane potential. Highly polarized mitochondria exhibit red color.

## Discussion

Neurological symptoms are a characteristic hallmark for type 2 and 3 GD. Unfortunately, no current therapies can prevent, slow, or halt GD’s neurodegenerative processes. ERT is not sufficient for treating neuropathology in GD because the recombinant enzyme does not cross BBB. Alternatively, small-molecules such as AMB that can cross the intact BBB and enhance GCase [26] or inhibit GC accumulation may overcome these limitations [27]. Two oral therapies, SRT and chaperone therapy, are designed to lower the accumulated GC in the lysosomes. If a chaperone therapy strategy is activated mutated GCase enzyme, the SRT approach is reduced GC production by inhibition of GluCer synthase. Our results show that the GluCer pathway inhibitor (EGT) and the pharmacologic chaperone (AMB) increase the residual enzyme activity in GD2 patients with L444P/L444P;Rec∆55;RecNCiI, and L444P/D409H mutations. However, AMB, but not EGT, increased GCase activity in cells with L444P/L444P;R495P/R495P;A456P mutations. Similar to AMB, EGT did not change the GCase activity in cells with L444P/L444P;D409H;A456P *GBA* mutation, as described before [15]. Our data fit previous reports that the AMB chaperone activity of not simply depends on the type of *GBA* mutation but seems to be patient dependent [12,15,28]. Other factors that can be involved in AMB pharmacodynamics may hypothetically surge efficiency; for example, increasing SapC and LIMP2 levels can stimulate AMB-induced GCase activity [20].

The cellular pathology of GD starts within the lysosome with chronic substrate accumulation. GC, as an integral part of glycosphingolipid metabolism, is involved in cell signaling transduction, membrane trafficking, or cytoskeletal processes [29]. Mutant GCase is recognized in cells as a misfolded protein. Instead of being trafficked to the lysosomes, it gets re-translocated to the cytoplasm, where it is degraded via the ubiquitin-proteasome system [28]. Several studies and our results confirm that AMB increases GCase trafficking to the lysosome and rescue the misfolded enzyme from degradation in GD macrophages and fibroblasts [28,30]. EGT reduces GC production by inhibition of transferase, UGCC.

However, little is known about the effects of EGT on the level and activity of GCase. The biochemical characteristic of EGT makes it unlikely that it is a lysosomal GCase inhibitor and thus may act as a PC [31]. Here, we report that EGT increased GCase activity in control, GD2, and GD3 cells. Based on data demonstrating that EGT enhanced GCase transport to the lysosomes, we suggest that improving trafficking and lysosomal pathway in the presence of EGT may explain how EGT increases GCase activity. In MPS fibroblasts, EGT was shown to correct the trafficking of lactosylceramide to the Golgi complex [32].

Dysfunctional lysosomes impair autophagy, block autophagic flux in GD [21,22,33], as demonstrated in multiple GD model systems, including GBA mouse models, GBA-/-flies, patient fibroblasts and PBMC, iPSC-neuronal model with GCase or saposin C deficiency [34-39]. Besides the EGT inhibitory activity and the chaperone activity of AMB, our results revealed that both molecules significantly induce autophagic flux, autophagosome-lysosomal fusion, and increased levels of acidic lysosomes in cells derived from GD2 and GD3 patients. As opposed to *GBA* variant dependent chaperone activity, the activation of autophagy-lysosomal processes in the presence of AMB and EGT is irrespective of *GBA* variants. Several studies have shown that AMB enhances lysosomal function in cellular models of Parkinson’s disease and reduces alpha-synuclein build-up, improving neuronal functions. Moreover, AMB triggers lysosomal exocytosis [40].

Interestingly, a study of EGT as a potential treatment for Parkinson’s disease also shows that EGT inhibits alpha-synuclein and stimulates autophagy flux in neurons by suppressing AKT-mTOR signaling in neurons [41]. Pharmacological induction of ALP can be a useful mechanism to promote GC clearance and protect cells against the secondary toxic effects. The induction of autophagy and reversal of lysosomal dysfunction may reduce neuronal cell death and potentially slow down neurologic disease progression in patients with GD2-3[42,43].

We found that AMB and EGT caused the significant inhibition of cell proliferation in GD2 and GD3 fibroblast. In some cases, the highest dose of AMB and EGT yielded a cytotoxic effect on GD cells. Previous *in vitro* studies indicated that 60 μM of AMB has a deleterious effect on wild-type mouse embryonic primary cortical neurons after 5 days of treatment [39], while the cause of decreased cell viability was not investigated. Some studies demonstrated that AMB did not cause cell death but disturbed mitochondrial membrane permeability [39]. EGT was shown to decrease the frequency of B cell malignancy in mice by the inhibition of UGCCwhich also slows the cell proliferation in liver cells [44,45].

Severe impairment of autophagy in GD2-3 leads to inhibition of mitophagy and mitochondrial metabolism [21]. However, suppression of autophagy and mitophagy in neural cells causes neurodegenerative progression [46]. Many studies have explored mitophagy and energy metabolism in GD2 and GD3 models [47,48]. However, only a few studies exhibited that AMB changed mitochondrial content in mouse neurons [39,49]. Cell viability assay indicates that both AMB and EGT induce total ATP production. The activation of mitochondrial functions by AMB and EGT demonstrated by increased mitochondrial mass and density, activation of mitochondrial membrane potential in GD2-3 fibroblasts indicates that there are similarities in the cellular response to AMB and EGT with regard to the type of *GBA* mutations.

## Conclusion

While substrate synthesis inhibition and pharmacologic chaperone therapies have different modes of action, their downstream effects improve lysosomal and mitochondrial functions.

## Supporting Information

**Supplemental Figure 1. Assessing the activity for AMB and EGT in PBMC and macrophages**: **A**. PBMC derived from healthy controls (n=5) and GD3 patients with L444P/L444P and L444P/R493C were cultured for 5 days in the presence of 100µM AMB and 10 µM EGT. Relative GCase enzyme activity was estimated as a fold change towards untreated control. Each bar represents the average +/-STDEV. ∗ p<0.05 compared with an untreated group. **B**. PBMC and macrophages derived from three GD3 patients with L444P/L444P mutation, as indicated in the figure, were treated for 5 days in the presence of AMB and EGT. Relative GCase enzyme activity was estimated as fold change towards untreated control. Each bar represents the average +/-STDEV. ∗ p<0.05 compared with an untreated group.

**Supplemental Figure 2. EGT and AMB increase autophagosome-lysosomal fusion in PBMC**. Autophagosome (green, DALGreen) and lysosome (red, LysoTracker) co-localization analysis in PBMC derived from healthy control and GD3 patients (P12, P13, and P14) with L444P/L444P *GBA* mutation.

**Supplemental Figure 3. Cell viability and metabolic status in individual GD2 and GD3 fibroblast cell lines. A**. P5 fibroblasts with L444P/L444P were treated with 1, 10, 100 µM of AMB and 0.1, 1,10 µM of EGT for 5 days. The CCK-8 cell viability assay, ATP content, and ATP/CCK-8 (cell viability) ratio were analyzed. The cell counting assay, CCK-8, and ATP results were normalized in relationship to the untreated cells. **B**. P6 fibroblasts with L444P/L444P;Rec∆55;Rec NCiI were treated with 1, 10, 100 µM of AMB and 0.1, 1,10 µM of EGT for 5 days. The CCK-8 cell viability assay, ATP content, and ATP/CCK-8 (cell viability) ratio were analyzed. The cell counting assay, CCK-8, and ATP results were normalized to the untreated cells. **C**. P7 fibroblasts with L444P/L444P;R495P/R495P;A456P *GBA* mutations were treated with 1, 10, 100 µM of AMB and 0.1, 1,10 µM of EGT for 5 days. The CCK-8 cell viability assay, ATP content, and ATP/CCK-8 ratio were analyzed. The cell counting assay, CCK-8, and ATP results were normalized to the untreated cells. **D**. P9 fibroblasts with L444P/D409H were treated with 1, 10, 100 µM of AMB and 0.1, 1,10 µM of EGT for 5 days. The CCK-8 assay, ATP content, and ATP/CCK-8 ratio were analyzed. The results were normalized to the untreated cells. Values are expressed as average ± SEM. ∗ p<0.05 compared with an untreated group.

**Supplemental Figure 4. Time course of cell viability in individual GD2 and GD3 fibroblast cell lines. A**. P5 fibroblasts with L444P/L444P were treated with AMB and EGT for 24, 48, 72 h, and 5 days. The CCK-8 was analyzed, and results were normalized to the untreated cells. **B**. P6 fibroblasts were treated with AMB and EGT for 24, 48, 72 h, and 5 days. The CCK-8 assay was measured, and results were normalized to the untreated cells. **C**. P7 fibroblasts were treated with AMB and EGT for 24, 48, 72 h, and 5 days. The CCK-8 assay was measured, and results were normalized to the untreated cells. Values are expressed as average ±SEM. ∗ p<0.05 compared with an untreated group.

**Supplemental Figure 5. Lactate dehydrogenase (LDH) assay**. Control and GD3 fibroblasts with L444P/L444P were treated with AMB EGT for 5 days. The LDH assay was analyzed, and results were normalized to the untreated cells. The data represents +/-SEM.

